# tRNA epitranscriptomic alterations associated with opioid-induced reward-seeking and long-term opioid withdrawal

**DOI:** 10.1101/2023.09.06.556560

**Authors:** J. Blaze, C. J. Browne, R. Futamura, V. Zachariou, E. J. Nestler, S. Akbarian

**Affiliations:** Department of Psychiatry, Icahn School of Medicine at Mount Sinai, New York, NY, USA; Department of Neuroscience, Icahn School of Medicine at Mount Sinai, New York, NY, USA; Friedman Brain Institute, Icahn School of Medicine at Mount Sinai, New York, NY, USA; Department of Pharmacology, Physiology, and Biophysics, Boston University Chobanian & Avedisian School of Medicine, Boston, MA USA; Department of Pharmacological Sciences, Icahn School of Medicine at Mount Sinai, New York, NY, USA; Department of Genetics and Genomic Sciences, Icahn School of Medicine at Mount Sinai, New York, NY, USA

## Abstract

DNA cytosine methylation has been documented as a potential epigenetic mechanism of transcriptional regulation underling opioid use disorder. However, methylation of RNA cytosine residues, which would drive another level of biological influence as an epitranscriptomic mechanism of gene and protein regulation has not been studied. Here, we probed whether chronic morphine exposure could alter tRNA cytosine methylation (m^5^C) and resulting expression levels in the medial prefrontal cortex (mPFC), a brain region crucial for reward processing and executive function that exhibits opioid-induced molecular restructuring. We identified dynamic changes in glycine tRNA (tRNA^Gly^_GCC_) cytosine methylation, corresponding to altered expression levels of this tRNA at multiple timepoints following 15 days of daily morphine. Strikingly, a robust increase in methylation, coupled with decreased expression, was present after 30 days of withdrawal, suggesting that repeated opioid administration produces changes to the tRNA regulome long after discontinuation. Furthermore, forebrain-wide knockout of neuronal *Nsun2*, a tRNA methyltransferase, was associated with disruption of reward-seeking in the conditioned place preference paradigm, and this effect was recapitulated by regional mPFC *Nsun2* knockout. Taken together, these studies provide a foundational link between the regulation of tRNA cytosine methylation and OUD and highlight the tRNA machinery as a potential therapeutic target in addiction.

## Introduction

Opioid use disorder (OUD) is a major continuing and worsening public health issue in the U.S., with a staggering 5.6 million people over the age of 12 diagnosed with OUD in 2021 alone^1^ and contributing to over 70,000 deaths per year due to opioid overdose^2^. While a large segment of the population continues to receive opioids for pain management, with the potential for misuse and addiction, it is crucial to identify preventative measures or therapeutic approaches for OUD. Many studies suggest that molecular changes to the brain’s reward circuit promote long-term functional regulation that underlies aberrant behavior in addiction^3^.

Considerable work has been performed to identify patterns of transcriptional regulation in the reward circuit to identify biological domains affected by drugs of abuse. However, most transcriptional changes are short-lived. Epigenetic mechanisms are thought to stamp in longer-term transcriptional regulation that drives persistent functional changes and behavioral patterns characteristic of addiction. Many studies have identified unique epigenetic transcriptional regulatory mechanisms in substance use disorders^4–6^, although much of this work has been focused on psychostimulants such as cocaine with less emphasis on opioids ^7^. Further, while most examined canonical histone marks and DNA methylation, very little attention has been paid to other forms of epigenetic regulation, specifically the role of epitranscriptomics.

Epitranscriptomics has emerged as a novel field bridging RNA expression and protein synthesis in neuronal function to outline how chemical modifications to RNA can dynamically impact translational processes^8^. These modifications are persistent, lending the ability to induce long-term dysregulation of neurobiological function which may serve as a largely unexplored mechanism mediating addiction vulnerability. While some studies have investigated drug-induced changes in m^6^A, a modification most prominent on mRNA^9,10^, no studies have explored the role of m^5^C, a chemical modification most prominent on transfer RNAs (tRNAs), in addiction. tRNAs are integral for protein synthesis, a process altered by opioid exposure ^11–13^, and m^5^C promotes tRNA stability in many mammalian tissues, including brain ^14–17^. Surprisingly, early studies of morphine administration showed changes to other realms of tRNA functioning (specifically, tRNA synthetase activity and amino acid binding to tRNAs)^18^ and our recent study identified numerous changes in expression of tRNA synthetase genes throughout the brain reward system following heroin self-administration in the mouse^4^. Changes to other aspects of tRNA function have also been explored in brain and peripheral tissues, including tRNA fragment abundance following exposure to opioids^19^ or other drugs of abuse^20, 21^. Taken together, these studies point to a potential role for changes in tRNA function in addiction, including OUD.

Our laboratory has shown that altering the tRNA epitranscriptome through deletion of the m^5^C writer protein NSUN2 in neurons alters neuroplasticity and behaviors related to neuropsychiatric disease via glycine-specific changes in tRNA expression, amino acid abundance, and translation efficiency^22^. Notably, many of these changes overlap with processes relevant to addiction, including memory and synaptic plasticity. Here, we probed whether changes to the tRNA epitranscriptome and the functionality of NSUN2 within the medial prefrontal cortex (mPFC), a brain region heavily implicated in opioid reward-processing and addiction pathogenesis^23, 24^, contribute to addiction-relevant behavior, and whether the observed opioid-induced changes persist through extended abstinence. To do this, we measured tRNA expression and m^5^C levels at previously identified candidate tRNAs^22^ in mPFC tissue from wild-type mice that received chronic morphine injections followed by an acute or extended withdrawal period. Further, we investigated opioid-seeking behavior in the morphine conditioned place preference test following knockout of neuronal NSUN2 in brain. Our findings point to the tRNA epitranscriptome as a novel candidate for molecular mechanisms driving OUD and open a new line of research targeting tRNA machinery as a potential therapeutic target in addiction.

## Methods

### Animals

All animal work was approved by the Institutional Animal Care and Use Committee of the Icahn School of Medicine at Mount Sinai. Mice were group-housed 2-5/cage with ad libitum access to food and water and a 12h light/dark cycle (lights off at 7pm) under constant conditions (21 ± 1°C; 60% humidity). Mice bred in-house were weaned at ∼P28, housed with same-sex littermates, and ear-tagged/genotyped.

### Nsun2 conditional knockout mice

*Nsun2* conditional knockout mice included forebrain *Nsun2* knockout (KO) (*Camk2a-Cre+,Nsun2^2lox/2lox^* vs. wildtype controls (*Camk2a-Cre-*)) and *Nsun2^2lox/2lox^* mice injected with either AAV8^hSyn1-CreGFP^ vs. AAV8^hSyn1-GFP^ as controls. Both conditional knockout models were previously validated in Blaze et al^22^. Mice with the mutant allele have exon 6 of *Nsun2* flanked by loxP sites, and presence of Cre-recombinase causes excision of exon 6 and a frame-shift resulting in early termination of *Nsun2* translation. Nsun2^2lox/2lox^ mice (2 copies of the mutant allele) were crossed with a *Camk2a-Cre+* line to produce *Camk2a-Cre+,Nsun2^2lox/2lox^* mice for knockout of Nsun2 in forebrain neurons, which is associated with widespread neuronal Cre-mediated deletion across the forebrain by the third postnatal week^25–28^. Age- and sex-matched *Camk2a-Cre-,Nsun2^2lox/2lox^* or *Nsun2^2lox/-^* were used as wild-type controls.

### Viral microinjections

For viral microinjection surgeries as described previously^22^, we injected virus into adult mouse mPFC (AAV8^hSyn1-CreGFP^ and AAV8^hSyn1-GFP^; see Fig. 1a for location and overlap with mPFC region of interest for rest of study), 10-12 week-old Nsun2^2lox/2lox^ mice were anesthetized with isoflurane and 1 μL of virus per hemisphere (bilateral injection) was injected at a rate of 0.25 μL/min using a Hamilton syringe (Reno, NV), a micropump (Stoelting) and a stereotactic frame (Stoelting). Stereotactic coordinates for mPFC injection were as follows: 1.5 mm anterior/ posterior, ±0.5 mm medial/lateral and 1.5 mm dorsal/ventral. Control animals received 1 μl per hemisphere of AAV8^hSyn1-GFP^ using the same conditions.

**Figure 1.**
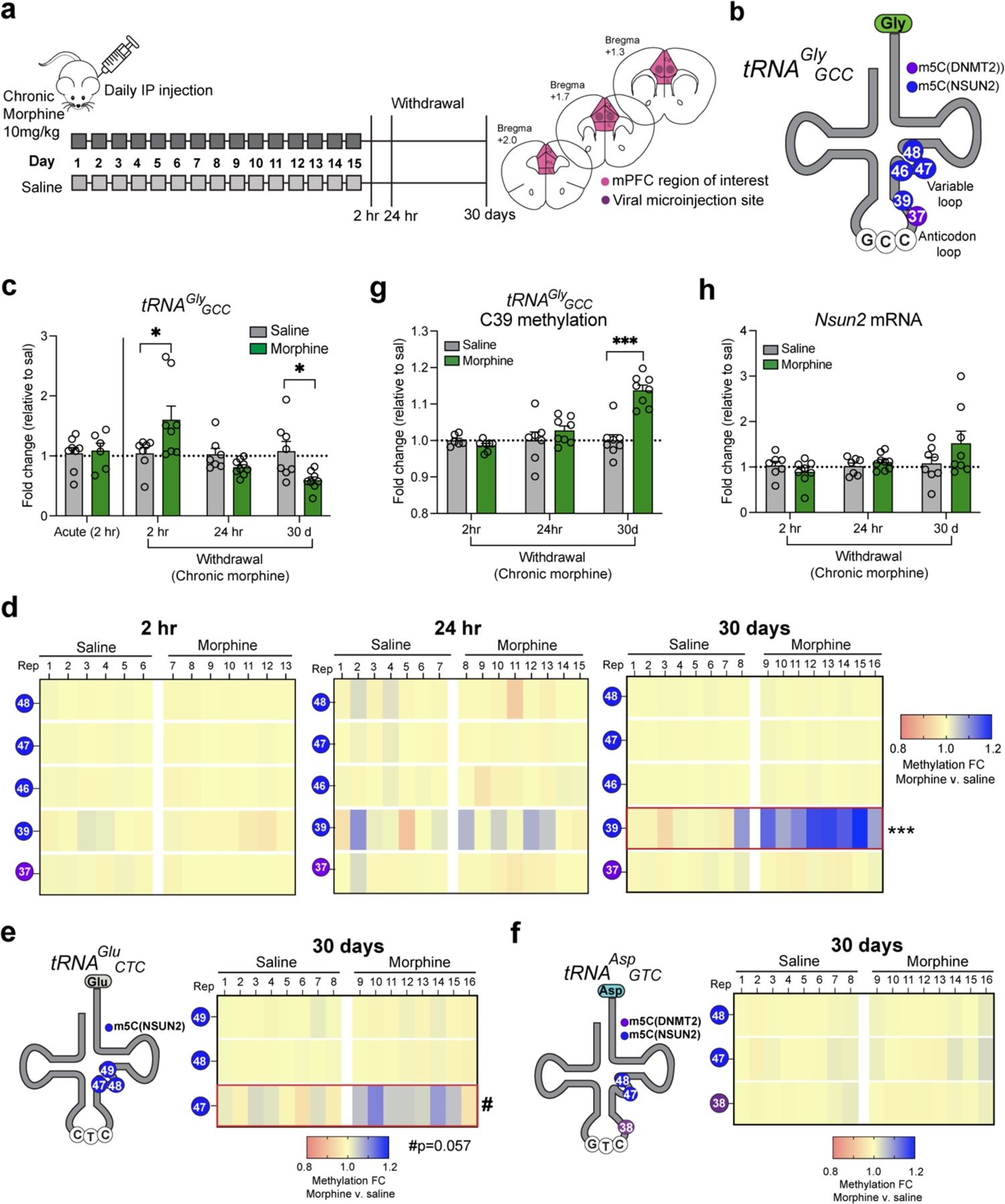
Dynamic tRNA^Gly^ expression and methylation following acute morphine and withdrawal in wild-type mice. **a)** Adult wild-type male mice were injected IP with 10mg/kg morphine daily for 15 days, followed by 2 hours, 24 hours, or 30 days of withdrawal and collection of medial prefrontal cortex (mPFC) tissue. See overlap of mPFC region of interest and viral Cre injections used in behavioral experiments **b)** Schematic of tRNA^Gly^GCC depicting methylated cytosines. **c)** qPCR for tRNA^Gly^GCC expression after a single acute morphine injection (n= 8 saline/6 morphine) or morphine withdrawal (2hr: n=7 saline/8 morphine, 24h: n=7 saline/9 morphine, 30d: n=8 saline/8 morphine; *p<0.05). **d)** Bisulfite amplicon sequencing for tRNA^Gly^GCC after 2hr, 24hr, and 30 days of withdrawal. Note the increase in methylation for morphine-treated mice at cytosine 39 at 30 days (t-test; ***p<0.001). **e)** Left, schematic representation of tRNA^Glu^CTC and cytosines methylated by Nsun2. Right, Bisulfite amplicon sequencing for tRNA^Glu^CTC showed a trending increase in methylation for morphine-treated mice at cytosine 47 (t-test #p<0.057; n=8/group). **f)** Left, schematic representation of tRNA^Asp^GTC and cytosines methylated by Nsun2 and Dnmt2. Right, Bisulfite amplicon sequencing for tRNA^Asp^GTC showed no changes in methylation at any sites between saline and morphine-treated mice (n=8/group). **g)** Cytosine 39 methylation at each timepoint shown as a fold change relative to the average of each group’s saline control (***p<0.001). **h)** qPCR for Nsun2 mRNA levels at all 3 withdrawal timepoints.

### Morphine Treatment

Morphine sulphate (from the National Institute on Drug Abuse) was diluted in saline and injected intraperitoneally at 10 mg/kg. For withdrawal studies, animals remained in the home cage for 2 hours, 24 hours, or 30 days after the last injection until sacrifice, at which point brains were extracted, and mPFC was dissected and flash frozen on dry ice.

### tRNA bisulfite sequencing

The tRNA amplicon bisulfite sequencing protocol was performed as described in Blaze et al.^22^ and previously adapted from Bormann et al.^29^. mPFC punches (see Fig. 1a) were homogenized in Trizol reagent (Invitrogen) and RNA was extracted, purified, and quantified using the Qubit fluorometer. 500ng of total RNA was used for bisulfite conversion with the EZ RNA Methylation kit according to manufacturer’s instructions (Zymo Research) but with an additional 2 PCR cycles to denature tRNAs completely, as previously reported^15^. cDNA synthesis was performed on bisulfite-treated RNA using the reverse primer for each of 3 tRNA isodecoders (i.e., *GlyGCC-1*^22, 30^*, GluCTC-1*^22^*, AspGTC-1*^22, 29, 30^) followed by PCR amplification with the tRNA-specific forward primer. Amplicons were size-verified on a 4% agarose gel and PCR amplicons were extracted and purified (Qiaquick Gel Extraction Kit, Qiagen) before indexing (Nextera XT, Illumina). and pooled up to 32 amplicons in equimolar concentrations before running with 75 bp paired-end reads on the MiSeq (Illumina).

### Data analysis

Bisulfite sequencing data was analyzed using BisAmp^29^, a publicly available web server for quantification of methylated reads from Miseq output fastq files. tRNA bisulfite sequencing data from bisAMP (methylation percentages for each cytosine) was further analyzed by conducting a t-test for each cytosine with FDR correction using the two-stage linear step-up procedure of Benjamini, Krieger and Yekutieli.

### YAMAT/UMI seq

Y-shaped Adapter MAture tRNA (YAMAT) sequencing was performed as previously described^22^ and was adapted from Shigematsu and colleagues^31^ to include unique molecular identifiers (UMIs) in order to avoid overamplification of individual tRNA isodecoders. Briefly, 5 µg total RNA was isolated from frontal cortex of morphine-or saline-treated animals. RNA was then deacylated to remove amino acids from 3’ ends and demethylated to remove m1A, m1C, and m3C modifications using a proprietary demethylation mix (Arraystar, Inc). 40 µm of YAMAT forked linkers were then incubated with pure demethylated and deacetylated RNA followed by addition of 10X annealing buffer (50 mM Tris HCl pH 8, 100 mM MgCl2, 5 mM EDTA) and then overnight incubation with T4 RNA ligase 2. Linker-ligated RNA was then incubated with RT Primer (TruSeq Small RNA Library Prep Kit, Illumina) and reverse transcription was performed with Superscript III RT (Invitrogen) followed by bead purification. Libraries were amplified using Phusion Hotstart II Polymerase (Thermo Scientific) with primers and indexes from the TruSeq Small RNA kit (Illumina) for 16 PCR cycles.

Amplified libraries were then bead-purified and run on the Agilent Bioanalyzer for confirmation of library size and quantified with the Qubit Fluorometer (Invitrogen). 8 libraries (n=4 morphine, 4 saline) were pooled at equimolar concentrations and run with 75 bp paired-end reads on the MiSeq (Illumina).

### Data analysis

Raw sequencing reads were processed using cutadapt to trim adaptor sequences and UMItools to extract UMIs from reads and perform deduplication. Paired end reads were merged using Pear and reads were filtered before differential expression analysis was performed using Deseq2, with significant differences in morphine vs. saline tRNA expression denoted at FDR-adjusted p<0.05.

### Real-time qPCR

For real-time qPCR to identify changes in *Nsun2* mRNA expression, we used 1 µg total RNA to generate cDNA followed by Taqman qPCR using Taqman Universal Master Mix and Taqman probes for *Nsun2* (Assay ID: Mm01349532_m1) and *Gapdh* as a control gene (Assay ID: Mm99999915_g1; Applied Biosystems). For qPCR to detect change in tRNA expression, we used the cDNA described above and pre-validated PCR primer sets specific to the mature tRNA sequence for *GlyGCC-1* or *5S rRNA* as a housekeeping gene (Arraystar, Inc,) using SYBR-based qPCR. Data were analyzed using the comparative Ct method and normalized to the housekeeping gene and saline-injected controls. An unpaired t-test was used to compare treatment groups and significance was denoted at p<0.05.

### Behavior

#### Morphine conditioned place preference (CPP)

CPP was carried out using an unbiased design in three-chamber Med Associates CPP boxes with overhead monitoring captured by a camera and analyzed using Ethovision (Noldus) to track the animal’s location within the box. The two end chambers differed in visual (stripe vs. gray) and tactile (small grid flooring vs. large grid flooring) cues to create distinct contexts. Dim white overhead light was used throughout testing. In the first session (pretraining), mice were placed in the CPP chamber and allowed to explore all three compartments freely for 20 minutes. Pairing sides were then determined for each mouse such that the group average of time spent was not biased for either chamber. Subsequently, mice received three days of morphine-context conditioning. On each conditioning day, mice were treated with saline and confined to one chamber for 45 min in the morning, and, in the afternoon, received morphine (10 mg/kg IP) and were confined to the other chamber for 45 min. On the fifth day, CPP was assessed by placing animals in the chamber and allowing them to explore freely. A place preference score was generated by subtracting the time the animal spent in the morphine-paired chamber vs. time spent in the saline-paired chamber.

#### Statistical analyses

Statistical analyses were performed using Graphpad Prism 8.4.3 software. Unpaired two-tailed t-tests were used for all comparisons, and FDR-correction was used for tRNA bisulfite amplicon sequencing. Statistical significance was denoted by p<0.05. All bar graphs are presented as the mean with error bars representing standard error of the mean (SEM) and include all individual data points.

## Results

### mPFC tRNA^Gly^ expression is dynamically altered during 30 days of morphine withdrawal

To assess whether a clinically relevant model of opioid addiction and withdrawal can alter tRNA expression and tRNA cytosine methylation in the rodent brain, wild-type adult male mice were injected intraperitoneally (IP) acutely (single dose) or daily for 15 consecutive days with 10 mg/kg morphine (or saline as control) and left undisturbed until sacrifice at one of three timepoints post-injection: 2 hours, 24 hours, or 30 days (**Fig. 1a**). The 30-day timepoint was chosen to coincide with a period of abstinence known to promote high levels of craving that drive relapse^32^, which we have shown to promote robust transcriptional regulation throughout the reward circuit^4^. In this well-validated opioid treatment paradigm, 10 mg/kg daily morphine has been shown to promote psychomotor sensitization and other reward- and addiction-related behavior, and many of these phenotypic changes have been linked to the mPFC, a crucial part of the well-known addiction circuitry in rodent models and human subjects^23, 24^. At each timepoint, mPFC, including infralimbic and prelimbic cortex (refer to **Fig. 1a**), was collected, and RNA was extracted to query tRNA levels and tRNA cytosine methylation following acute or chronic morphine treatment or withdrawal.

As a starting point, we chose to quantify tRNA^Gly^ levels because this tRNA family is highly sensitive to alterations in tRNA cytosine methylation. This sensitivity is likely due to the fact that brain is one of the organs with the highest levels of tRNA cytosine methylation, altogether encompassing 5 cytosine methylation sites^22^, the most of any tRNA family and not seen for any of the other tRNAs assigned to the other 23 canonical amino acids. Specifically, of the four tRNA^Gly^ isoacceptors (ACC, CCC, GCC, TCC), tRNA^Gly^_GCC_ (**Fig. 1b**) is the most highly expressed in brain and sensitive to disruptions in cytosine methylation^22^. Thus, we assessed tRNA^Gly^_GCC_ expression in the mPFC of morphine- and saline-treated mice by qPCR. Indeed, there was a dynamic change in tRNA^Gly^_GCC_ expression levels throughout the 30 day withdrawal period (2-way ANOVA treatment x timepoint interaction; F(3,53)=6.118, p=0.001; acute: n= 8 saline/6 morphine, 2hr: n=7 saline/8 morphine, 24h: n=7 saline/9 morphine, 30d: n=8 saline/8 morphine), defined by a transient increase followed by a significant decline in the long-term. Specifically, tRNA^Gly^_GCC_ was increased 1.6-fold in morphine-treated mice 2 hours after the last injection (t(53)=3.063, p=0.014; n=7 saline/8 morphine) but showed no change at 24 hours (t(53)=1.215, p=0.919; n=7 saline/9 morphine). Conversely, after 30 days of withdrawal, morphine-treated mice showed a 0.6-fold decrease in tRNA^Gly^_GCC_ expression (t(53)=2.756, p=0.032; n=8 saline/8 morphine) (**Fig. 1c**), suggesting a dynamic pattern of tRNA^Gly^_GCC_ expression throughout the withdrawal period after repeated morphine exposures. In contrast, no alterations were observed after a single dose of morphine (t(53)=0.243, p>0.999; **Fig. 1c**), suggesting that dynamic changes in tRNA^Gly^_GCC_ expression are driven by chronic opioid exposure.

Because tRNA^Gly^_GCC_ showed a significant decrease in morphine-treated animals after 30 days of withdrawal in chronic morphine-exposed animals, we asked whether other tRNA^Gly^ isoacceptors show similar types of alterations. We quantified tRNA^Gly^_TCC_ and observed a marginal, non-significant decrease in mPFC expression levels after 30 days of morphine withdrawal (**Supp. Fig. 1a-b**; unpaired two-tailed t-test, t(13)=1.398, p=0.186; n=8 saline, 7 morphine), suggesting that tRNA^Gly^ may have isoacceptor-specific regulation following repeated exposure to opioids and long-term withdrawal. Next, we examined whether these effects of morphine withdrawal on tRNA^Gly^ expression were specific to the infralimbic and prelimbic regions of the mPFC (which are firmly implicated in opioid addiction circuitry) or alternatively representative of changes broadly affecting the entire rostral cortex **(Supp.** Fig. 1a,c**,d)**. Therefore, we repeated the morphine exposure and 30-days withdrawal paradigm in a different cohort of mice and dissected out the entire rostral cortex (see **Supp.** Fig. 1c for exact location). There were no differences between treatment groups for tRNA^Gly^_GCC_ levels by qPCR (t(18)=1.136, p=0.271; n=10 saline/10 morphine) or by YAMATseq (**Supp. Fig. 1e**; n=4 saline/4 morphine), an unbiased screen of the tRNAome using next-generation sequencing (see Methods). Notably, the input required for YAMATseq was prohibitive for conducting this analysis on mPFC specifically. We conclude that morphine withdrawal-induced changes in tRNA^Gly^_GCC_ are specific to the mPFC without affecting other areas of rostral cortex.

### tRNA methylation in mPFC is increased following 30 days of morphine withdrawal

Having shown that withdrawal from opioids affects tRNA^Gly^_GCC_ expression, we then wondered if a mechanism for these types of fluctuation in tRNA levels are associated with tRNA epitranscriptomic dysregulation, given that previous studies in peripheral cell lines have uncovered a strong link between tRNA expression levels and cytosine methylation changes following stress ^14, 15^. Because we have previously shown decreased methylation is directly linked to decreased tRNA^Gly^_GCC_ expression and our data here point to dynamic alterations in tRNA^Gly^_GCC_ in mPFC after 2 hours and 30 days of morphine withdrawal, we queried the methylation status of tRNA^Gly^_GCC_ in mPFC of saline- and morphine-injected mice at all three morphine withdrawal timepoints. We used tRNA bisulfite sequencing with approximately 40,000 reads per sample to measure tRNA^Gly^_GCC_ cytosine methylation levels in mPFC of mice that experienced morphine withdrawal (or saline) at 2 hr, 24 hr, and 30 days and identified a significant 1.15-fold increase in methylation only after 30 days for morphine-treated mice at cytosine 39 (t(14)=6.330, FDR adj. p<0.0001; n=8 saline/8 morphine), while methylation at all other cytosines remained unaltered by morphine exposure (**Fig. 1d,g; Supp. Table 1**).

Notably, methylation at C39 (and other cytosines) were unchanged by morphine treatment at earlier timepoints of 2 hours and 24 hours into withdrawal, showing a temporal specificity for methylation changes that only appear after protracted abstinence **(Fig. 1d,g; Supp. Table 1**). Because we saw a significant increase in tRNA^Gly^_GCC_ methylation at 30 days, we also measured cytosine methylation at other tRNA families that contain fewer cytosine methylation sites. First, tRNA^Glu^ contains 3 methyl-cytosines (**Fig 1e**), and bisulfite sequencing of tRNA^Glu^_CTC_ revealed a near significant methylation increase at cytosine 47 for morphine-treated mice after 30 days (**Fig 1e**; t(14)=2.654, FDR adj. p=0.057; n=8 saline/8 morphine; **Supp. Table 1**). We also examined tRNA^Asp^_GTC_ which likewise contains 3 methyl-cytosines and found no morphine-induced changes after 30 days of withdrawal (**Fig 1f**; **Supp. Table 1**).

In brain, tRNA cytosine methylation is mainly deposited by NSUN2, which methylates ∼80% of tRNAs, while the other RNA methyltransferase, DNMT2, methylates a much smaller subset of tRNAs. Among the methylcytosines within tRNA^Gly^_GCC_, 4 out of 5 are methylated by NSUN2 (C39, C46-48, **Fig. 1b**), while one site is DNMT2-dependent (C37). Conversely, tRNA^Glu^_CTC_ has only 3 NSUN2-dependent methylation sites (C47-49; **Fig 1e**), and tRNA^Asp^_GTC_ carries 2 NSUN2-mediated cytosines and 1 DNMT2-mediated cytosine (**Fig. 1f**). We noticed that the subset of cytosines altered after morphine withdrawal (tRNA^Gly^_GCC_ C39 and tRNA^Glu^_CTC_ C47) are both methylated by NSUN2, suggesting that tRNAs with more NSUN2-dependent cytosine methylation sites may be selectively vulnerable to changes associated with long-term withdrawal from opioids. Therefore, we quantified NSUN2 levels by qPCR at each withdrawal timepoint in mPFC and found a positive correlation between methylation levels at tRNA^Gly^_GCC_ cytosine 39 (the only cytosine modified by morphine withdrawal) and *Nsun2* mRNA levels only for the morphine-treated animals (**Fig. 2a**; saline: r^2^=0.014, p=0.600, n=22 mice, 7-8 per timepoint; morphine: r^2^=0.234, p=0.017, n=24 mice, 7-8 per timepoint), thereby suggesting that dynamic changes in morphine-induced tRNA hypermethylation could, at least in part, be linked to corresponding adaptations in *Nsun2* expression. However, we did not detect significant changes in *Nsun2* mRNA levels between morphine and saline-treated animals in the mPFC at 2 hr, 24h, or 30 days **(Fig. 1g-h**; 2-way ANOVA, n=7-8 saline/8-9 morphine; all p’s>0.05). Additionally, tRNA^Gly^_GCC_ cytosine 39 methylation levels negatively correlated with tRNA^Gly^_GCC_ expression levels only in morphine-treated animals (**Fig. 2b**; r^2^=0.307, p=0.005, n=24 mice, 7-8 per timepoint) while no correlation was observed in saline-treated animals (**Fig. 2b**; r^2^=0.093, p=0.179, n=24 mice, 7-8 per timepoint). Taken together, our data showing increased tRNA^Gly^_GCC_ cytosine methylation potentially driven by NSUN2 coupled with decreased tRNA^Gly^_GCC_ expression suggest that long term morphine withdrawal induces long-term changes in the mPFC tRNA regulome, including NSUN2-mediated tRNA cytosine methylation and expression.

**Figure 2.**
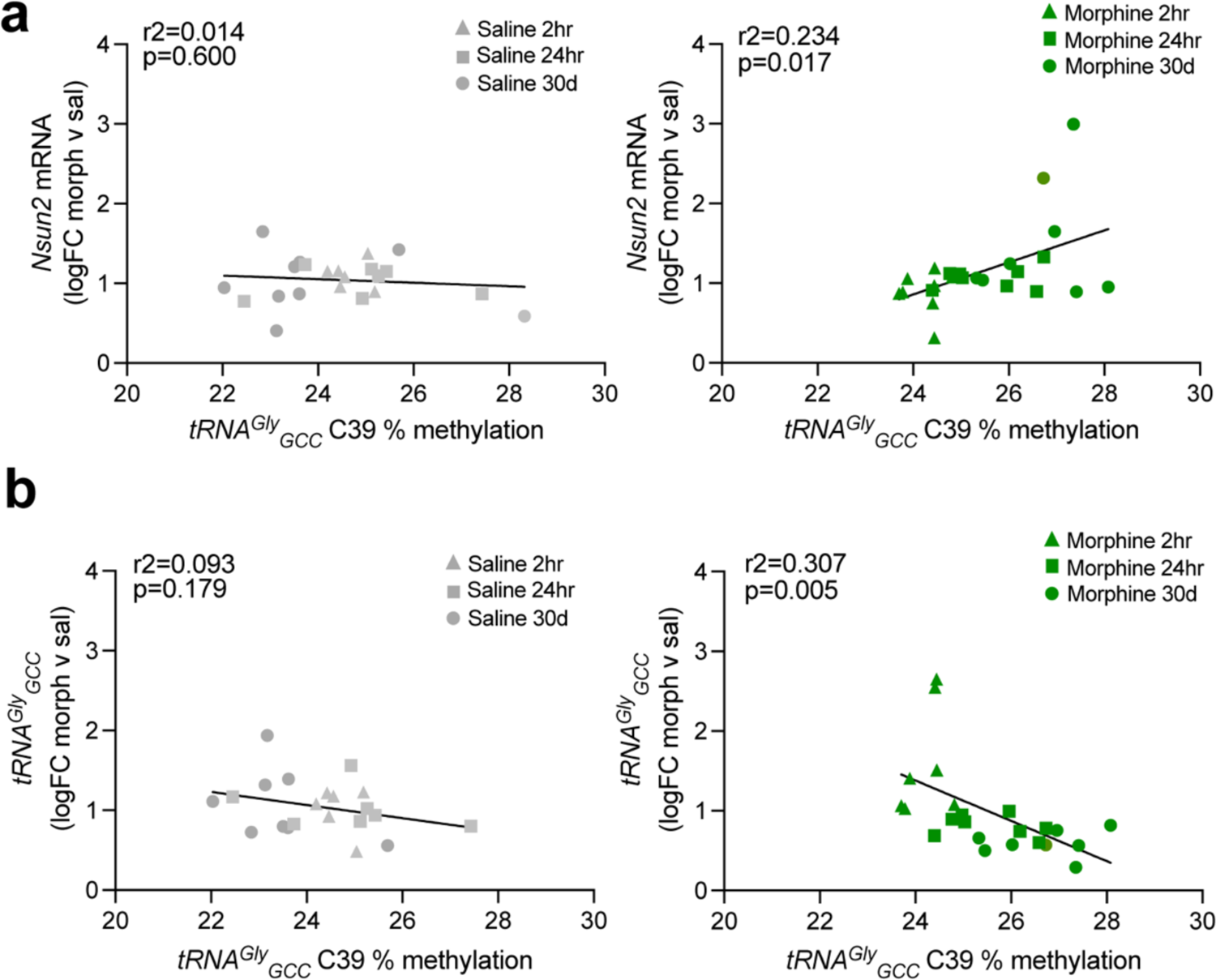
Correlations between tRNA^Gly^GCC methylation and **a)** Nsun2 mRNA levels and **b)** tRNA^Gly^GCC expression levels. Left, saline groups show no significant correlation or anticorrelation, while the morphine groups (right) show significant relationships between expression levels of tRNA and Nsun2 and C39 methylation.

### Neuronal Nsun2 deletion impairs opioid reward seeking

Given our findings linking NSUN2-regulated tRNA^Gly^_GCC_ methylation and expression following long-term morphine withdrawal in the mPFC, we then tested whether deletion of *Nsun2* impacts addiction-related behavior. Early studies revealed alterations in protein synthesis in synaptosomal fractions following acute and chronic morphine exposure^11, 12^, suggesting that protein translation in neurons, and potentially regulatory molecules such as tRNAs in neurons, are involved in opioid addiction phenotypes. Therefore, we generated mice with conditional neuron-specific knockout of *Nsun2* in forebrain (*Camk2a-Cre+, Nsun2^2lox/2lox^* mutant mice) driven by *Ca^2+^*/*calmodulin-kinase II (CamK)-Cre* promoter, which produces widespread neuronal *Cre*-mediated deletion across the forebrain before postnatal day 18 as previously described. We conducted a standard morphine CPP paradigm using the same dose as the aforementioned long-term morphine exposure paradigm (Fig. 1a), which is a well-established task to measure reward seeking and has translational relevance to the drug-environment associations in human OUD^33^. Intact drug-cue interactions and reward seeking in CPP require expression of addiction-related proteins, but it is unknown whether dysfunction of the neuronal tRNA regulome is involved in CPP performance. To test this, *Nsun2 Camk2a-Cre* conditional knockout (cKO) mice and wild-type controls underwent a traditional morphine CPP task (see Methods; **Fig. 3a**). As expected, wild-type mice showed a highly significant preference for the morphine-paired side of the chamber (one-sample t-test, t(6)=5.037, p=0.002, n=7 WT), whereas in striking contrast *Nsun2 Camk2a-Cre* cKO mice as a group completely lacked preference over baseline (one-sample t-test, t(7)=0.3191, p=0.759, n=8 KO) and spent significantly less time in the morphine-paired side compared to wild-type controls (unpaired t-test, t(13)=2.194, p=0.047; n=7 WT, 8 KO; **Fig. 3b**).

**Figure 3.**
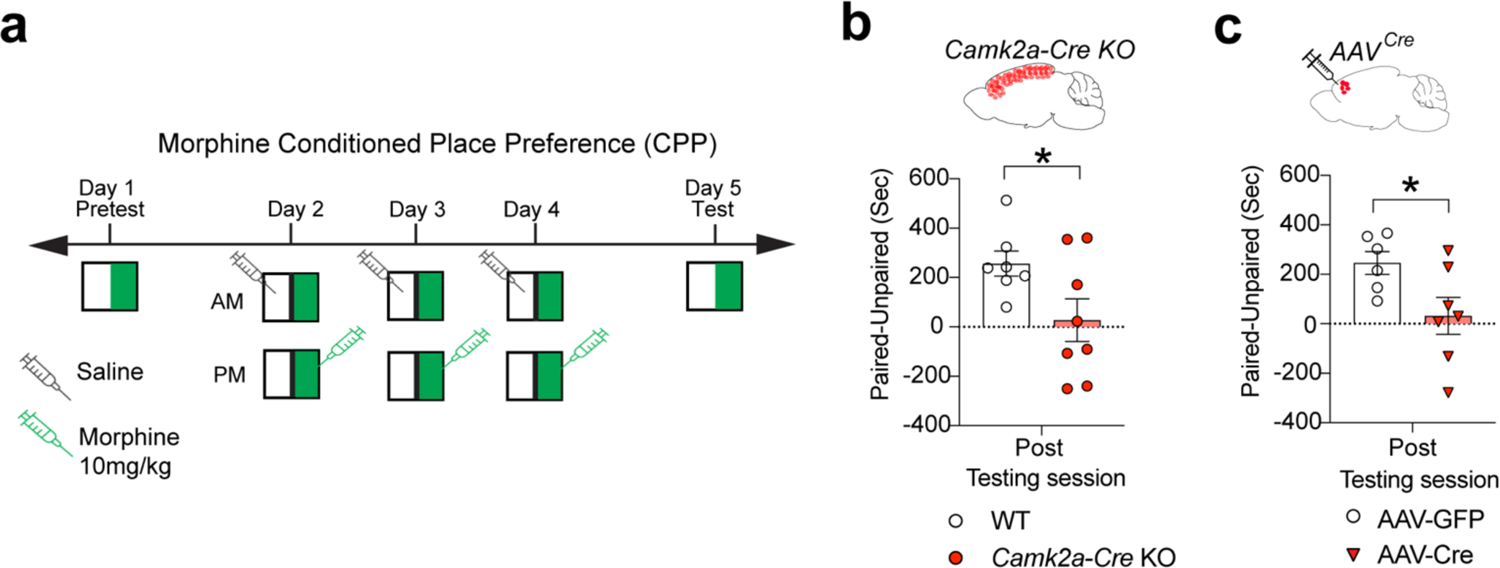
Morphine conditioned place preference in Nsun2 conditional knockout mice. **a)** For CPP, mice received a 20-min pre-test wherein all compartments of the chamber could be explored. Subsequently, mice underwent three 45-min conditioning sessions in which saline was injected in the morning restricted to one side of the chamber (“unpaired”), and 10 mg/kg morphine was injected in the afternoon on the other side (“paired”). Conditioning to morphine was assessed in a subsequent 20-min session with access to both sides of the chamber, with positive conditioning indicated by a higher preference on the morphine-paired side. **b)** Camk2a-Cre Nsun2 KO mice (n=7 WT, 8 KO) and **c)** PFC AAV-Cre Nsun2 KO (n=6 AAV-GFP, 7 AAV-Cre KO) mice both show a reduction in time spent in the morphine-paired chamber compared to the unpaired chamber, indicating a blunted reward response to morphine (t-test *p<0.05).

Because forebrain deletion of *Nsun2* may affect numerous psychological and behavioral processes that broadly impair function, we examined whether the observed impairment in opioid reward seeking after *Nsun2* knockout could be recapitulated by specifically targeting the mPFC. To test this, we conducted the same type of CPP experiment in 12–16-week-old conditional *Nsun2^2lox/2lox^* mice that received a highly localized bilateral mPFC microinjection of AAV8^hSyn1-CreGFP^ (AAV-Cre), or of AAV8^hSyn1-GFP^ as a control (AAV-GFP) for region- and cell-type-specific knockout of neuronal *Nsun2* (see **Fig. 1a** for overlap with mPFC region of interest used in molecular studies). Previous studies have shown that this area of mPFC is essential for opioid-induced reward behavior but not for contextual memory. Deletion of *Nsun2* in adult neurons of the PFC produced a similar impairment in morphine CPP (**Fig. 3c**; AAV-GFP vs. baseline one sample t-test, t(6)=5.332, p=0.003; AAV-Cre vs. baseline one sample t-test, t(7)=0.429, p=0.683; AAV-GFP vs. AAV-Cre unpaired t-test, t(11)=2.337, p=0.039; n=6 AAV-GFP/7 AAV-Cre), suggesting that mPFC *Nsun2* expression, and possibly NSUN2-mediated tRNA methylation, in this discrete brain region is necessary for opioid-reward seeking behavior.

## Discussion

We show here that chronic morphine exposure elicits dynamic changes in expression of tRNA^Gly^_GCC_ in the rodent mPFC, characterized by an early increase in expression and ultimately leading to a decrease in expression in morphine-treated animals after 30 days of withdrawal. Furthermore, changes in expression are tightly linked to site-specific NSUN2-mediated cytosine methylation levels at tRNA^Gly^_GCC_. We also demonstrate that loss of neuronal NSUN2 in forebrain or more locally within a discrete area of the mPFC impairs opioid-seeking behavior, suggesting the importance of the m^5^C epitranscriptomic system in regulating addiction phenotypes. As the first investigation of tRNA cytosine methylation in the context of addiction, we now provide a previously unexplored regulatory system, the tRNA epitranscriptome, as an important mediator of opioid action.

There is a wealth of evidence that chronic exposure to opioids induces dynamic and long-term changes to transcriptional machinery via epigenetic modifications to chromatin in the brain’s reward circuitry^4-7^, but very few studies have investigated the epitranscriptome in relation to addiction. To our knowledge, the only investigation into CNS epitranscriptomic changes in relation to opioids focused on m^6^A, showing that chronic morphine administration altered m^6^A profiles in spinal cord for many circular RNAs (circRNAs), non-coding RNAs that are single-stranded and covalently bound in a closed loop, which may be a molecular change contributing to the effects of morphine tolerance^10^. The m^6^A epitranscriptome was also implicated in alcohol use disorder (AUD), with a small cohort of human postmortem brains showing AUD-specific patterns of methylation at various mRNAs, long non-coding RNAs, and miRNAs^9^. The current study is the first to investigate the m^5^C tRNA epitranscriptomic in the context of addiction. Because tRNAs are crucial components of protein translation in the ribosome, and many studies have pointed to protein synthesis abnormalities in the brains of morphine-exposed rodents ^11–13^, it is surprising that there has been such little investigation into tRNA regulatory mechanisms in opioid addiction.

While we profiled cytosine methylation and full-length tRNA expression, there are reports of drug-induced changes to tRNA fragments of varying sizes originating from various tRNA isoacceptor families. In the rat nucleus accumbens, tRNA fragments were altered following methamphetamine self-administration, including a decrease in the fragment originating from the 5’ end of tRNA^Gly^_GCC_ ^21^. Taken together with our findings of NSUN2 and cytosine methylation correlations in brain following opioid exposure, the results suggest that the tRNA regulome has a drug- and tissue-specificity that may contribute differentially to addiction phenotypes.

Furthermore, there is evidence that tRNA-mediated effects of drugs of abuse are not limited to the nervous system. For example, in the male reproductive system, including sperm and epidydimosomes, which are tissues that have been well-characterized as carrying large amounts of tRNA fragments that are sensitive to stressful conditions including low-protein diet ^34^. Furthermore, chronic ethanol exposure in rodents altered NSUN2 expression and increased the abundance of tRNA fragments originating from tRNA^Glu^_CTC_ and tRNA^His^_GTG_ in the male reproductive tract, while fragments from tRNA^Ser^_AGA_ were decreased in sperm^20^. Conversely, another study measured tRNA fragments in sperm of human opioid users and found a decrease in various tRNA fragments, with the most marked effect being at tRNA^Gly^_GCC._^19^.

Importantly, changes in NSUN2-mediated tRNA methylation in the mature brain selectively decrease mature tRNAs ^22^. For this reason, in the present study we chose to focus on mature tRNA expression and m^5^C in the brains of morphine-exposed mice, but future studies could link our findings with tRNA^Gly^_GCC_ fragmentation. The downstream molecular or behavioral sequalae of these changes have not been elucidated. In peripheral tissues and cultured cells, it has been established that impairments to the tRNA methylation machinery via ablation of NSUN2 causes accumulation of tRNA fragments, which activates cellular stress pathways and globally decreases protein synthesis^14, 15^.

Here we identified 30 days after morphine withdrawal a significant morphine-dependent increase in tRNA^Gly^_GCC_ C39 methylation that correlated with an increase in tRNA^Gly^_GCC_ expression. The question remains: How can an increase in tRNA expression modulate the addiction circuitry to confer vulnerability to future drug-seeking? To our knowledge, this is the first evidence of a site-specific experience-dependent increase in methylation of m^5^C, as most studies demonstrate loss of methylation following environmental perturbations ^14, 15, 35^. Therefore, investigating how increased methylation drives tRNA abundance is unknown. Our group has previously shown that viral overexpression of NSUN2 in wild-type mouse mPFC selectively increases methylation at tRNA^Gly^_GCC_ C39 and also confers a depressive-like phenotype ^22^. While this provides an additional link between increased tRNA^Gly^_GCC_ C39 methylation and downstream behavioral consequences, future studies will identify if virally overexpressing *Nsun2* also increases tRNA^Gly^_GCC_ abundance and how this contributes to depressive-like behavior, which involves overlapping circuitry as that involved in addiction.

tRNAs can sense environmental cues and modulate the epitranscriptomic landscape in an experience-dependent manner. While this has mostly been demonstrated in the context of toxin exposure^35^ or cellular stress via UV radiation^14, 15^, there is evidence of global tRNA m^1^A levels increasing following non-associative learning in the cerebral ganglia of *Aplysia californica* ^36^. We now add to this concept by showing drug- and withdrawal-dependent dynamic changes in the tRNA m^5^C epitranscriptome in mammalian brain. Not only are these observed molecular changes dynamic, but the presence of a robust tRNA phenotype at 30 days after the last morphine injection provides new insight into mechanisms by which the brain may be primed for relapse or reinstatement after prolonged abstinence. Therefore, investigation of the tRNA regulome during relapse or reinstatement is of interest as a future direction of this work.

## Acknowledgments

We thank Dr. Andrew Chess for providing sequencing equipment. This work was supported by NIH grant F32MH115565 to JB, R01DA054526 to SA, R01NS086444 to VZ, and P01DA047233 and R01DA007359 to EJN.

## Author Contributions

J.B. and C.J.B. conceptualized and designed experiments. C.J.B. and R.F. performed mouse morphine dosing and CPP behavior experiments and analyzed CPP behavioral data. J.B. performed all viral surgeries for CPP, conducted molecular assays, and analyzed and interpreted all molecular data. S.A., E.J.N., and V.Z. contributed design of experiments and interpretation of data. J.B., C.J.B, and S.A. wrote the manuscript with input from all authors.

---

**Supplementary Figure 1.**
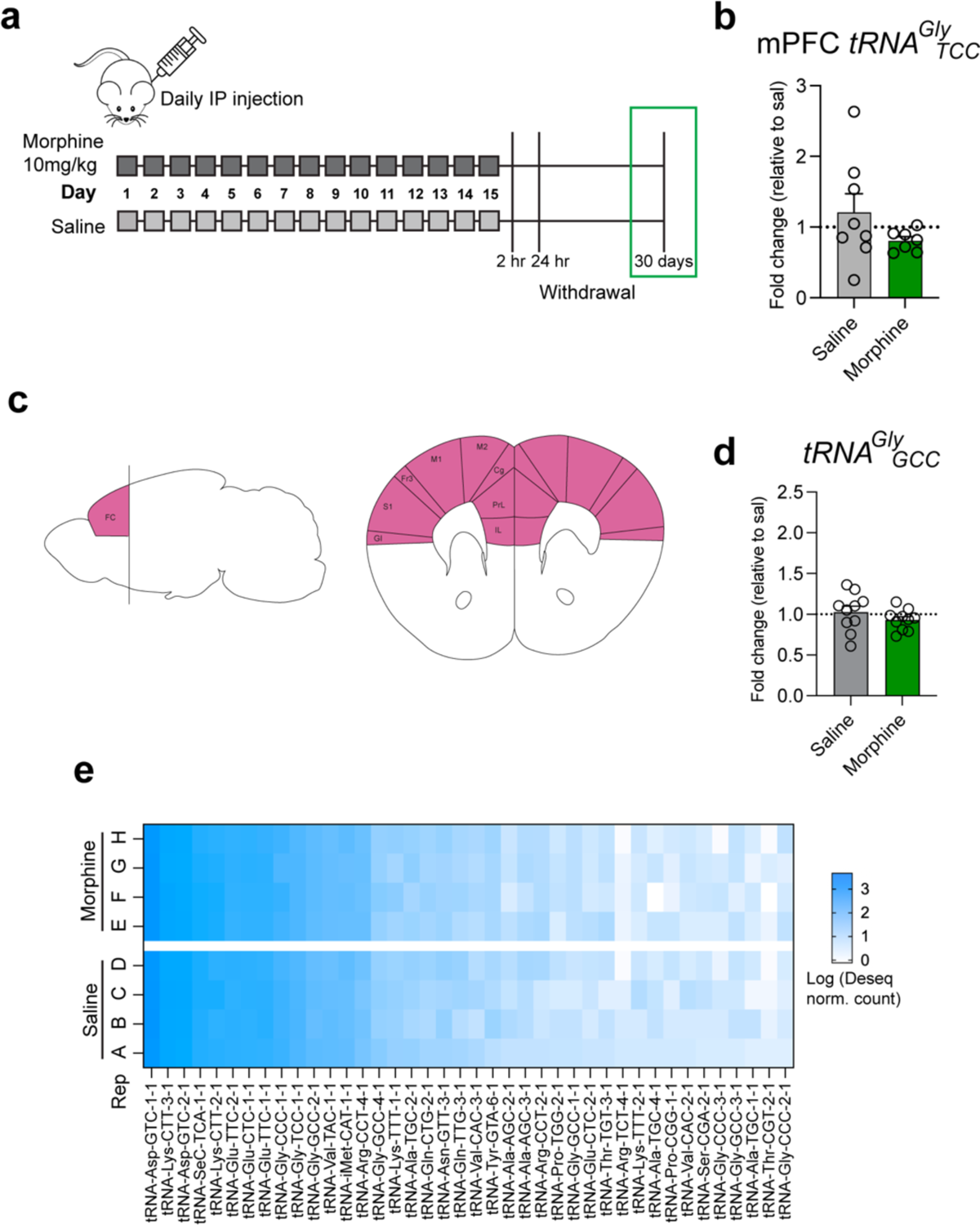
**a)** After 30 days of withdrawal, mPFC was collected and showed no change in expression of another tRNA^Gly^ isoacceptor, tRNA^Gly^TCC. **b)** Brain atlas localization of tissue taken for frontal cortex molecular assays. **d)** After 30 days withdrawal, frontal cortex showed no change in tRNA^Gly^GCC expression (n=10/group). **e)** Heatmap shows top 40 isodecoders that were most highly and reliably expressed in all samples using Deseq normalized counts.

**Supplementary Table 1.**
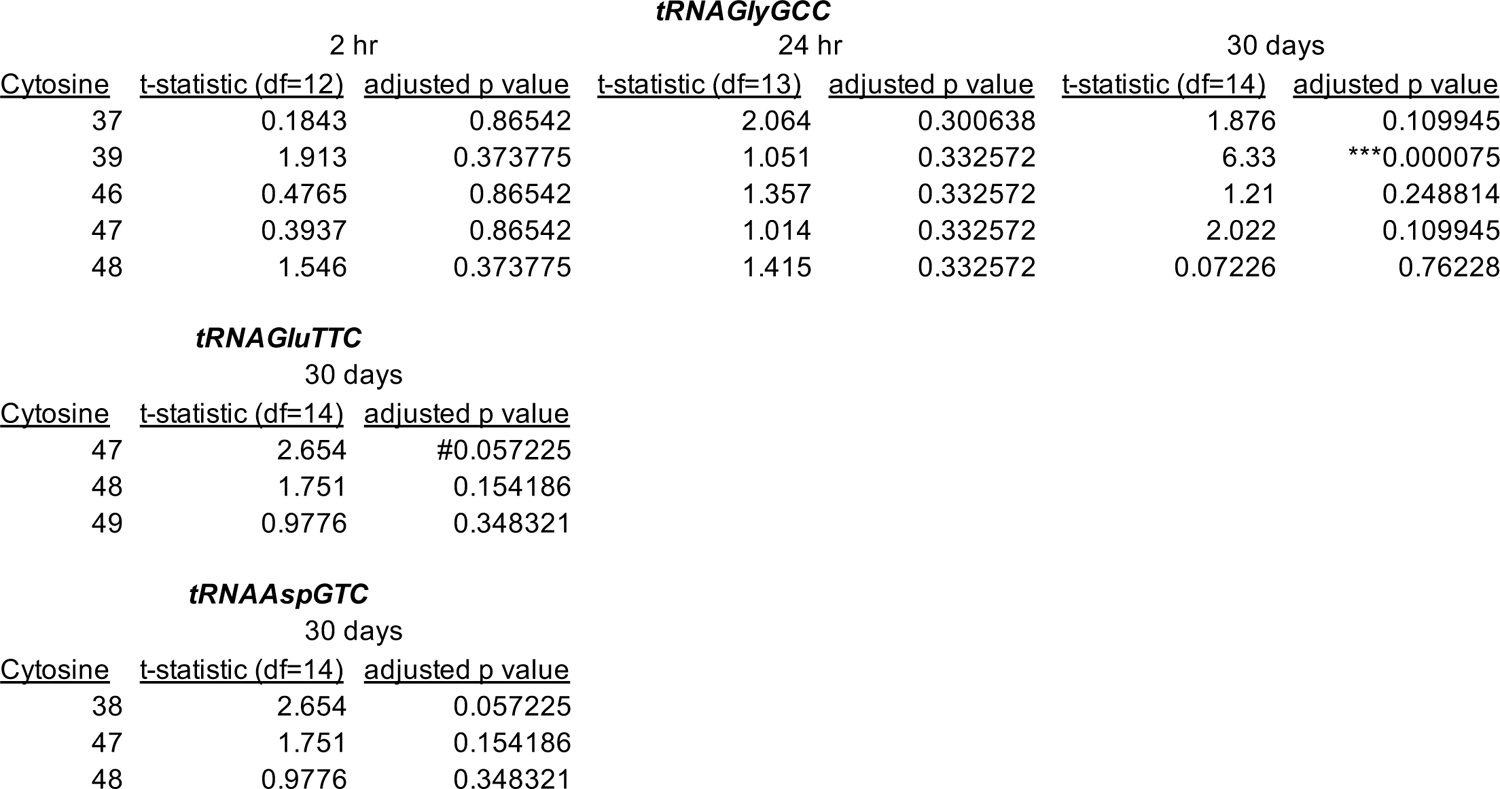
Site-specific statistical analysis of bisulfite tRNA sequencing.

